# Mirror-Image L-DNA Nanocubes for Stable and Targeted Multimodal Drug Delivery

**DOI:** 10.64898/2026.05.20.726533

**Authors:** Terry Farkaly, Sihan Wu, Yuliya Dantsu, Arifuzzaman Tapash, Wen Zhang

**Affiliations:** Department of Biochemistry, Molecular Biology and Pharmacology, Indiana University School of Medicine, 635 Barnhill Drive, Indianapolis, IN 46202; Department of Chemistry, Indiana University, Bloomington, Indiana 47405, United States; Melvin and Bren Simon Comprehensive Cancer Center, 535 Barnhill Drive, Indianapolis, IN 46202, USA

**Keywords:** mirror-image DNA, nanocube, chemotherapy, siRNA, folic acid

## Abstract

Nucleic acid nanostructures provide programmable architectures for molecular delivery but remain limited by rapid nuclease degradation, poor in vivo persistence and inefficient intracellular cargo release. Here we report a mirror-image L-DNA nanocube as a biologically persistent and modular therapeutic delivery platform. The nanocube self-assembles from synthetic L-DNA oligonucleotides into a structurally defined architecture that exhibits substantially enhanced resistance to enzymatic degradation and prolonged stability under physiological conditions compared with the corresponding D-DNA nanostructure. Surface functionalization with folic acid enables selective tumour targeting in vitro and in vivo. The L-DNA nanocube supports the delivery of chemically distinct therapeutic cargos, including doxorubicin, a bortezomib prodrug and MCL1-targeting small interfering RNA (siRNA). In tumour-bearing mice, L-DNA nanocube-mediated delivery improves therapeutic efficacy while reducing systemic toxicity relative to free drug and D-DNA nanocube controls. For siRNA delivery, we engineer a pH-responsive release mechanism that promotes endosomal escape and cytosolic cargo localization, as visualized by cryo-electron tomography, resulting in efficient gene silencing. Together, these results establish mirror-image nucleic acid nanostructures as a class of biologically functional nanomaterials for programmable intracellular therapeutic delivery.

## INTRODUCTION

Nucleic acid nanotechnology enables the programmable assembly of structurally precise architectures with applications in biosensing, molecular computation, therapeutics and nanomedicine^1, 2^. Sequence-defined nucleic acid interactions provide exceptional control over nanoscale geometry, molecular organization and cargo positioning, allowing the construction of dynamic nanostructures with tunable physicochemical and biological properties^3^. Over the past two decades, advances in DNA and RNA self-assembly have generated increasingly sophisticated nanostructures for molecular recognition, intracellular delivery and therapeutic modulation^4^. Despite these advances, the translational potential of nucleic acid nanomaterials remains substantially constrained by their limited stability and functionality in biological environments. A major challenge for nucleic acid nanostructures is their susceptibility to rapid degradation by endogenous nucleases and their poor persistence under physiological conditions^5^. Structural destabilization during circulation can compromise cargo retention, targeting efficiency and intracellular delivery, thereby limiting therapeutic performance in vivo^6, 7^. Numerous approaches have been explored to improve the biological stability of nucleic acid nanostructures^8^, including backbone modification^9^, polymer conjugation^10^, lipid encapsulation^11^ and inorganic surface coating^12^. Although these strategies can partially protect nucleic acid assemblies from degradation, they may give rise to unanticipated toxicities^13, 14^, reduce structural programmability or interfere with the intrinsic addressability of nucleic acid architectures^15^. Furthermore, many delivery systems remain vulnerable to inefficient intracellular trafficking and endosomal sequestration, particularly for therapeutic nucleic acids that require cytosolic access for activity.

Biological systems exhibit strong molecular chirality, and naturally evolved nucleic acid-processing enzymes preferentially recognize D-nucleic acids^16^. Mirror-image nucleic acids composed of L-ribose or L-deoxyribose therefore represent a fundamentally distinct class of synthetic genetic polymers that are intrinsically resistant to endogenous nucleases and largely orthogonal to native nucleic acid-processing pathways (Fig. 1A)^17–22^. Previous studies have demonstrated the utility of L-nucleic acids in molecular recognition and therapeutic aptamer development, including Spiegelmers with enhanced serum stability and pharmacological persistence^23, 24^. More recently, mirror-image biology has emerged as a broader framework for engineering synthetic biomolecular systems with altered interactions with living organisms, including the use of L-nucleoside molecules as antiviral and antitumor agents^22^, L-RNA aptamers to bind to disease targets^25–28^, L-DNA tetrahedron structures as drug delivery vehicles^29^, pH-responsive i-motif structure formed by L-DNA for therapeutic use^30^, and L-DNA hydrogels, as stable 3D cell-culture matrices and bio-scaffolds^31^. However, the construction of structurally defined mirror-image nucleic acid nanomaterials for programmable intracellular therapeutic delivery remains largely unexplored.

**Figure 1.**
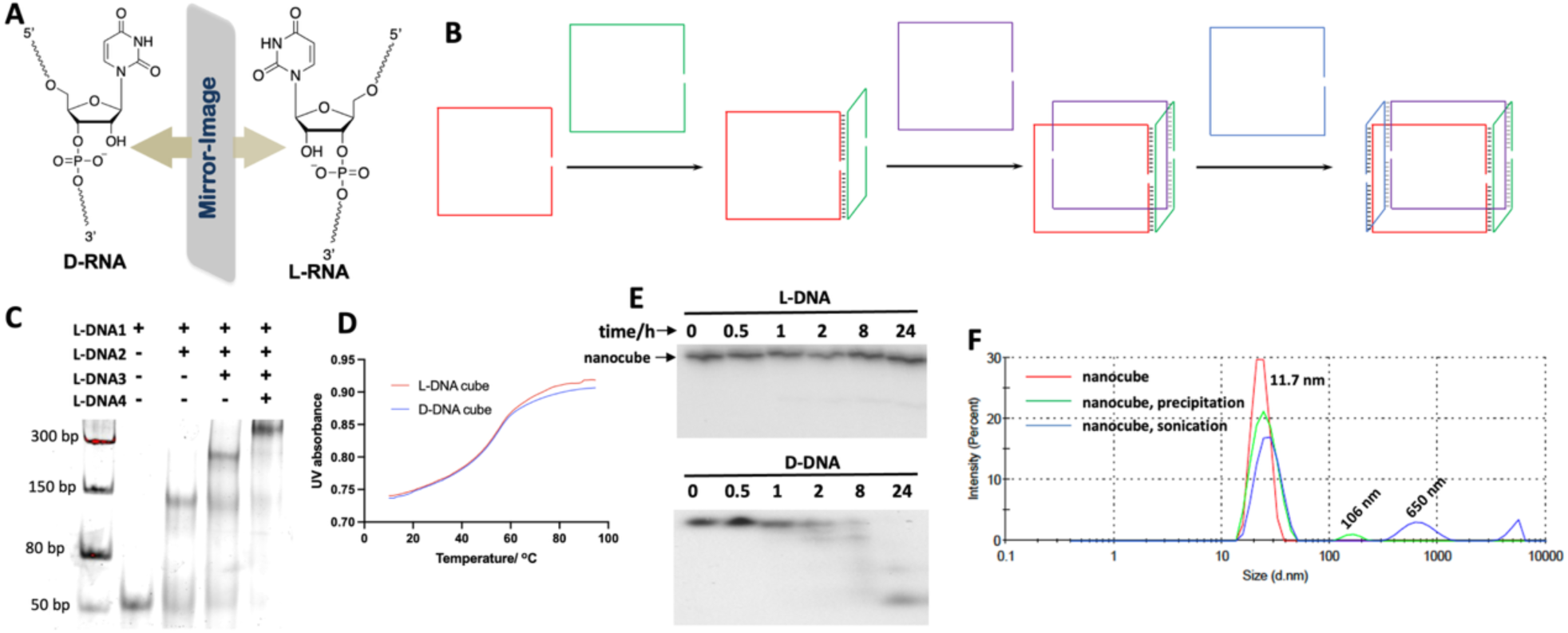
Design, Construction and Characterization of L-DNA nanocube. (A) Chemical structures of D- and L- ribonucleotides; (B) Schematic illustration of the self-assembled L-DNA nanocube architecture composed of four complementary L-DNA oligonucleotides. (C) native PAGE showing the assembly of the nanocube, stained by SYBR gold. (D) *T*_m_ melting curves for the assembly of the L-and D-nanocube. (E) Stability assay of L-nanocube (top) and D-nanocube (bottom) after treating with serum. (F) Dynamic light scattering (DLS) analysis of L-DNA nanocubes under native conditions and following ethanol precipitation or sonication treatment.

In addition to extracellular stability, intracellular delivery presents a critical barrier for nucleic acid therapeutics^32^. Efficient cytosolic delivery requires coordinated control over cellular uptake, endosomal trafficking and cargo release. Many nanocarriers exhibit substantial endosomal trapping, leading to limited bioavailability of therapeutic payloads in the cytoplasm^33^. Although a variety of endosome-disrupting materials and responsive delivery systems have been developed, achieving programmable intracellular release while maintaining structural precision remains challenging for nucleic acid nanostructures. Direct visualization of intracellular cargo localization and endosomal escape also remains technically difficult, limiting mechanistic understanding of nucleic acid nanocarrier function in living cells.

Here we report a mirror-image L-DNA nanocube that functions as a biologically persistent and modular platform for intracellular therapeutic delivery. The nanocube self-assembles from synthetic L-DNA oligonucleotides into a structurally defined architecture with substantially enhanced nuclease resistance and physiological stability relative to corresponding D-DNA nanostructures. Through programmable surface engineering, folic acid-functionalized L-DNA nanocubes enable selective tumour targeting in vitro and in vivo. The platform supports the delivery of chemically diverse therapeutic cargos, including doxorubicin, a bortezomib prodrug and MCL1-targeting small interfering RNA (siRNA), resulting in improved therapeutic efficacy and reduced systemic toxicity in tumour-bearing mice. For siRNA delivery, we further engineer a pH-responsive release mechanism that promotes endosomal escape and cytosolic cargo localization, visualized by cryo-electron tomography, leading to efficient gene silencing. Together, our findings establish mirror-image nucleic acid nanostructures as biologically functional nanomaterials for programmable intracellular therapeutic delivery and expand the translational scope of nucleic acid nanotechnology.

## MATERIALS AND METHODS

### Preparation of L-DNA and chimeric oligonucleotides, and functional L-DNA nanocube structures

The L-DNA, and chimeric oligonucleotides used for biochemical, cellular and in vivo studies were solid-phase synthesized using ABI 394 DNA/RNA synthesizer. The phosphoramidite compounds are from Chemgenes or Glen Research, Inc. The DMTr-on in-house synthesized oligonucleotides were deprotected by AMA (1:1 mixture of ammonium hydroxide and methyl amine solution) for 15 min at 65°C, followed by the Et_3_N•3HF treatment; and purified by Agilent ZORBAX Eclipse-XDB C18 column using 25 mM triethylammonium acetate in H_2_O (pH 7.5) with an increasing gradient of 0 % to 30 % acetonitrile over 40 mins. The 5’-Cyanine 5 labelled oligonucleotides were deprotected by conc. ammonium hydroxide at room temperature for 36 hours, followed by the Et_3_N•3HF treatment and denaturing urea polyacrylamide gel electrophoresis. To label the L-DNA with folic acid (FA), the alkyne modified L-DNA was synthesized using the 1-Ethynyl-dSpacer CE Phosphoramidite and 2’-alkyne modified phosphoramidites from Hongene Biotech, Inc., following the standard deprotection protocol. Long oligonucleotides (>50 nt) were purified by denaturing PAGE. After purification, the oligonucleotides were desalted and re-concentrated to the proper concentrations for nanostructure construction and biochemical experiments.

All the sequence information and the MS characterization of L-nucleic acids are listed in Supporting Information. All the nanostructures were self-assembled by mixing the different strands at the same molar concentrations in TMS buffer (25 mM Tris-HCl pH 7.5, 10 mM MgCl_2_, 50 mM NaCl) or PBS buffer (137 mM NaCl, 2.7 mM KCl, 10 mM Na_2_HPO_4_, 2 mM KH_2_PO_4_, pH 7.4), heated to 90°C for 2 min and slowly cooled to 23 °C. After the annealing program, the nanostructures were loaded onto 10% native PAGE, stained by Sybr Gold (Invitrogen), and visualized by Bio-rad ChemiDoc Imaging System.

### Click reaction to label L-DNA with folic acid molecule and peptide

To facilitate the specific cellular targeting and the escape of the oligonucleotides from endosome after receptor-mediated endocytosis, the L-DNA was synthesized with alkyne modification, which later was conjugated with a folic acid molecule via a click reaction. The folic acid-PEG-azido (MW 1kDa) compound was purchased from Biopharm PEG, Inc. The azido labelled INF7 peptide was purchased from Aapptec LLC. To the alkyne modified L-DNA solution, excess of folic acid-PEG-azide or peptide-PEG-azide, tris(3-hydroxypropyltriazolylmethyl)amine/CuSO_4_ and sodium ascorbate solution were added. The reaction continued for 30 min, followed by ethanol precipitation and gel purification, to get folic acid or peptide labelled L-DNA. Details of the click reaction procedure are included in the Supporting Information.

### L-DNA enzymatic stability assays

Both L- and D-3WJ nanostructures were incubated in 25% Human serum off-the-clot (Sigma) at 37°C for different times. 200 ng of RNA were taken at 30 min, 1 hr, 2 hr, 8 hr and 24 hr time points, and analyzed by 10% native PAGE in TB buffer. The gels were stained with Sybr Gold (Invitrogen) and washed briefly before imaging using Bio-rad ChemiDoc Imaging System.

### Melting temperature measurement of L-DNA nanocube

Melting temperature measurement was performed using the Varian Cary 300 UV-Visible Spectrophotometer equipped with Agilent temperature controller. The experiments were performed using the samples (0.2 µM RNA or DNA structures) dissolved in the buffer of 50 mM NaCl, 10 mM NaH_2_PO_4_-Na_2_HPO_4_ (pH 6.5), 0.1 mM EDTA and 10 mM MgCl_2_. The samples were heated to 90 °C and allowed to cool down to room temperature slowly. The UV absorbance at 260 nm was recorded at a heating rate of 0.5 °C/min.

### Cell culture

MDA-MB-468 (RRID: CVCL_0419) and HEK-293T (RRID: CVCL_0063) cells were purchased from the American Tissue Type Collection (ATCC, Manassas, VA), or gift from Indiana University Melvin and Bren Simon Comprehensive Cancer Center. Cells were maintained in DMEM (Corning, Cat. #10-013-CV) supplemented with 10% FBS. All cell lines were maintained at 37°C in a fully humidified atmosphere containing 5% CO_2_, used at early passage for experiments, and tested to be free of mycoplasma contamination. To evaluate the targeting and drug delivery efficiency of the L-DNA nanostructures conjugated with folic acid, cell culture experiments were conducted in a folic acid-depleted environment. Cells were grown in folic acid-free medium (SigmaAldrich, catalog # D2429) for 16 hours prior to introducing the L-DNA-folic acid conjugate. This step ensured that folic acid from external sources was minimized, reducing background interference and allowing the effects of the folic acid-L-DNA nanostructure conjugate to be more accurately attributed to receptor-mediated uptake.

### Confocal microscopy imaging of internalization of L-DNA nanocube

Cells were grown on poly-d-lysine-coated glass cover slides in normal growth media as described above and allowed to attach overnight. Cells were grown in folic acid-free medium (SigmaAldrich, catalog # D2429) for 16 hours prior to the experiments. For confocal microscopy, cells were fixed by 4% paraformaldehyde, permeabilized using PBS-T (0.01% Triton X) and stained by DAPI for nuclei visualization. FITC-conjugated phalloidin is used to stain the cell cytoskeleton. Confocal microscopy was performed using a Zeiss AxioObserverZ1 modified by 3i for confocal microscopy with the addition of a CSU-X1 M1 Spinning Disk Confocal and a Prime BSI CMOS Detector.

### Flow cytometry analysis

Flow cytometry was performed to quantify the attachment of Cy5⁺, FA⁺ and Cy5⁺, FA⁻ L-DNA nanocubes. MDA-MB-468 cells were cultured in DMEM with 10% FBS at 37°C in a 5% CO₂ incubator until they reached 80% confluence. Cells were seeded in 6-well plates at a density of 1 × 10⁵ cells per well and allowed to adhere overnight. Nanostructures were added at a final concentration of 50 nM in folic acid-free medium and incubated with cells for 4 hours. After incubation, cells were washed three times with PBS, detached and resuspended in 1 mL PBS for analysis. Flow cytometry was performed with Cy5 fluorescence detected and at least 10,000 events were collected for each sample. Data were analyzed using FlowJo software (FlowJo LLC), and the data are presented as the percentage of the cells.

### Dynamic light scattering

The hydrodynamic diameter of L-DNA nanocubes was measured by dynamic light scattering (DLS) in Physical Biochemistry Instrumentation Facility, IU Bloomington. Samples were prepared under native aqueous conditions and analysed using a DLS instrument according to the manufacturer’s instructions. For comparison, aliquots subjected to ethanol precipitation or sonication were analysed under the same measurement conditions. The size distribution was recorded as intensity-weighted particle diameter, and the resulting profiles were used to assess nanocube monodispersity and aggregation behaviour.

### Formation of L-DNA nanocube/Doxorubicin complexes

Doxorubicin hydrochloride (HPLC-purified, Sigma-Aldrich, catalog # D1515) was dissolved in Milli-Q water for a 2 mM stock solution, divided into aliquots and stored at −20°C. The Doxorubicin loaded L-DNA nanocube was formed by mixing the pre-assembled nanocube with DOX at 1:5 ratio for 24 h. The DOX-loaded nanocube was further conjugated with a folic acid labelled L-DNA for cellular experiments with the corresponding concentrations.

### Alamar blue assay for cell toxicity measurement

The Alamar Blue assay is based on the reduction of resazurin, a non-toxic and cell-permeable dye, to resorufin, a fluorescent compound, by metabolically active cells^34^. This reduction can be quantified via fluorescence or absorbance, providing a measure of cell viability and proliferation. For Alamar blue assay, cells were seeded into poly-d-lysine-coated 96-well plates at 8000-10000 cells per well in normal growth media as described above and allowed to attach overnight. Different drugs were diluted in folic acid-free medium to appropriate concentrations before the treatment. Cells were grown in folic acid-free medium (SigmaAldrich, catalog # D2429) for 16 hours prior to the treatment with different drugs. After treatment, Alamar Blue reagent (Invitrogen, Cat. #A50101) was added to each well, with the final volume of 10% in the well. The cells with Alamar Blue were further incubated at 37°C for 6 hours, and the fluorescence intensity (excitation 560 nm, emission 590 nm) was measured using Synergy H1 microplate reader (BioTek Instruments, Inc., VA). The fluorescence readings were analysed to determine cell viability, and the data were plotted.

### In vivo fluorescence imaging

For in vivo biodistribution and tumour-targeting studies, fluorescently labelled L-DNA nanocubes, corresponding D-DNA nanocube, free DOX and PBS buffer were administered to tumour-bearing mice by tail vein injection. In vivo fluorescence imaging was performed using an IVIS Spectrum imaging system at WuXi Biologics LLC. MDA-MB-468 tumour xenografts were established in NSG mice, and treatment was initiated when tumours reached approximately 80 mm^3^. Nanocube formulations were prepared in sterile phosphate-buffered saline (PBS) and injected intravenously in a total volume of 200 μL per mouse. Mice received three injections administered every other day over a 6-day treatment period. Mice were anesthetized with isoflurane (3–4% induction and 1–2% maintenance in oxygen) and maintained on a heated imaging stage throughout image acquisition. Fluorescence images were collected at 2, 6, 24, 48 and 72 h after the final injection to evaluate nanoparticle biodistribution and tumour accumulation. Quantitative fluorescence analysis was performed using Living Image software (PerkinElmer). Control and treatment groups were imaged under identical acquisition conditions for direct comparison of fluorescence intensity and tissue localization profiles.

### Animal studies

Female NSG or NRG mice (4–8 weeks old) were maintained under specific pathogen-free conditions in the Laboratory Animal Resource Center at Indiana University School of Medicine. All animal experiments were performed in accordance with protocols approved by the Institutional Animal Care and Use Committee (IACUC) of Indiana University School of Medicine (Protocol #25028). For tumour xenograft studies, MDA-MB-468 human breast cancer cells (1 × 10^7^ cells in 100 μL PBS) were orthotopically injected into the mammary fat pad. Tumour growth was monitored twice weekly using caliper measurements, and tumour volume was calculated using the formula V = (length × width²)/2. Treatment was initiated when tumours reached approximately 80 mm^3^.

Mice were randomly assigned to treatment groups receiving DOX/L-DNA nanocube formulations, DOX/D-DNA nanocube controls, free doxorubicin or vehicle (PBS). Nanoparticle formulations and free drug controls were administered by tail vein injection every other day for a total of three injections over 6 days. Tumour growth and body weight were monitored throughout the study. Animals were euthanized 14 days after treatment initiation or earlier if humane endpoint criteria were reached, including tumour volume exceeding 1000 mm^3^, body weight loss greater than 20%, ulceration or impaired mobility.

### Therapeutic efficacy and toxicity evaluation

Therapeutic efficacy was evaluated by monitoring tumour growth and body weight throughout the treatment period. At the experimental endpoint, mice were euthanized and blood samples were collected for plasma biochemical analysis. Plasma urea levels were measured using a commercial colorimetric assay kit according to the manufacturer’s instructions. Haematoxylin and eosin (H&E) staining was done by WuXi Biologics LLC, including major organs of liver, kidney, heart, lung and spleen. The organs were harvested, fixed in formalin, embedded in paraffin and sectioned for haematoxylin and eosin staining.

### Confocal colocalization analysis of intracellular trafficking of siRNA

Cells were seeded onto glass-bottom dishes and incubated with fluorescently labeled siRNA-loaded nanocubes. After treatment, cells were washed with phosphate-buffered saline (PBS) twice. For endocytic trafficking analysis, cells were fixed with paraformaldehyde, permeabilized with Triton X-100, and blocked with bovine serum albumin. Samples were then stained with primary antibodies against clathrin or RAB7, followed by incubation with appropriate green fluorescent secondary antibodies. Nuclei were counterstained with DAPI. Confocal fluorescence images were acquired using a laser scanning confocal microscope under identical acquisition settings across groups. Colocalization between siRNA and endocytic markers was assessed. Representative images were collected from three independent experiments.

### Immunogold transmission electron microscopy (TEM)

To visualize intracellular localization of delivered siRNA, biotinylated siRNA cargo was incorporated into L-DNA nanocube formulations and detected using streptavidin-conjugated nanogold particles (Nanopartz Inc.). Cancer cells were incubated with siRNA-loaded nanocubes, washed with PBS, and chemically fixed for electron microscopy analysis.

For optimized immunogold detection, post-embedding labeling procedures were performed without osmium tetroxide (OsO_4_) treatment to improve accessibility of biotinylated RNA to streptavidin-conjugated nanogold particles. Embedded sections were incubated with streptavidin-conjugated nanogold at optimized dilution conditions, followed by transmission electron microscopy imaging. Control samples were processed in parallel under identical conditions, by treating MDA-MB-468 cells with PBS buffer.

### Synthesis of azide-functionalized bortezomib boronate ester prodrugs and their conjugation to L-DNA

To enable covalent conjugation of bortezomib onto the L-DNA nanocube platform, azide-functionalized bortezomib boronate ester prodrugs were synthesized through modification of the boronic acid pharmacophore. Three structurally distinct boronate ester derivatives were prepared using different boronate ester architectures designed to modulate hydrolytic stability and intracellular release properties. Each prodrug incorporated an azide functional group to permit subsequent click conjugation to alkyne-modified oligonucleotides. Bortezomib boronic acid was converted into the corresponding boronate ester intermediates using the indicated diol or linker components under standard organic synthesis conditions. Products were purified by chromatography and characterized by NMR spectroscopy and mass spectrometry.

L-DNA nanocubes containing alkyne-functionalized oligonucleotide handles were first assembled through programmable sequence hybridization and annealing. Following nanocube assembly, azide-functionalized bortezomib boronate ester prodrugs were conjugated to the pre-assembled nanocubes using copper-catalysed azide–alkyne cycloaddition (CuAAC) chemistry^35^. This post-assembly click functionalization strategy enabled covalent installation of bortezomib prodrugs onto intact mirror-image nanocube scaffolds while preserving nanostructure integrity. Following click conjugation, excess unreacted small-molecule prodrugs and reaction components were removed using size exclusion chromatography.

### Intracellular activation and cytotoxicity analysis of bortezomib-loaded nanocubes

Following receptor-mediated endocytosis, intracellular acidification within endosomal compartments was expected to promote hydrolysis of the boronate ester linkage, thereby regenerating active bortezomib intracellularly. To evaluate biological activity, cancer cells were treated with free bortezomib, empty nanocube controls or bortezomib-loaded L-DNA nanocubes containing the different prodrug structures. Cell viability was measured after treatment using Alamer Blue assay. Dose-dependent effects were analysed across multiple concentrations of each bortezomib prodrug formulation.

## RESULTS AND DISCUSSIONS

### Design, construction and characterization of L-DNA nanocube

To establish a biologically persistent mirror-image nucleic acid nanostructure for therapeutic delivery, we designed a self-assembled L-DNA nanocube composed of four synthetic L-DNA oligonucleotides programmed through complementary base-pairing interactions (Fig. 1B). The nanocube architecture was designed to preserve the structural programmability of conventional DNA nanotechnology while introducing complete mirror-image chirality through replacement of natural D-deoxyribose with L-deoxyribose. Each strand contained defined hybridization domains that directed formation of a closed three-dimensional nanocube structure under aqueous annealing conditions. Assembly of the L-DNA nanocube was confirmed by native gel electrophoresis, which revealed progressive mobility shifts corresponding to stepwise strand incorporation and formation of the fully assembled nanostructure (Fig. 1C).

### Stability of L-DNA nanocube

To evaluate the biophysical stability of the mirror-image nanostructure, we next compared the thermal and enzymatic properties of L-DNA and corresponding D-DNA nanocubes. Melting temperature analysis demonstrated stable hybridization of the L-DNA architecture, consistent with efficient Watson–Crick base pairing within the mirror-image scaffold (Fig. 1D). The melting temperature of L-nanocube was measured to be 59.2 °C, which is comparable to the melting temperature of D-nanocube (T_m_ is 58.8 °C). It is beneficial for future therapeutic applications at physiological temperatures. We next tested the stability of the L-DNA nanostructures in human serum. Both L- and D-nanocubes were incubated with 25% human serum for up to 24 hours and the presence or absence of degradation products were assessed by gel electrophoresis (Figure 1E). The band of the intact L-DNA nanostructure remained stable throughout all time points, while the native DNA was completely degraded within 1 hour. These data indicate the dramatically enhanced resistance of L-DNA nanocube to nuclease degradation relative to the native D-DNA nanocube when incubated under biologically relevant conditions.

These findings establish that mirror-image L-DNA oligonucleotides can self-assemble into structurally defined and biologically stable three-dimensional nanostructures while preserving the programmability of canonical nucleic acid nanotechnology. The combination of nanoscale structural precision, colloidal stability under physiological conditions and exceptional nuclease resistance supports the use of L-DNA nanocubes as biologically persistent delivery scaffolds for intracellular therapeutic applications.

### Dynamic light scattering analysis reveals aggregation of L-DNA nanocubes following dehydration and mechanical perturbation

To evaluate the physicochemical properties of the assembled mirror-image nanostructures, the hydrodynamic size distribution of the L-DNA nanocube was analysed by dynamic light scattering (DLS) (Fig. 1F). Under native aqueous conditions, the nanocube exhibited a dominant particle population centred at approximately 11.7 nm, consistent with the expected dimensions of the programmed nanostructure and supporting successful assembly into a discrete nanoscale architecture.

In contrast, substantial changes in particle size distribution were observed following ethanol precipitation or sonication treatment. Ethanol precipitation induced the appearance of significantly larger particle populations, including aggregates exceeding 100 nm in diameter, whereas sonication further promoted the formation of micron-scale assemblies. These observations indicate that both dehydration-associated processing and mechanical perturbation can destabilize colloidal dispersion of the nanocube and promote interparticle aggregation. The aggregation behaviour observed is likely attributable to disruption of hydration-mediated stabilization and altered electrostatic interactions between nucleic acid nanostructures during solvent exchange. And sonication may introduce local mechanical stress to the cubic structure that facilitate intermolecular association between partially perturbed nanocubes.

Our findings provide several important insights into the physicochemical behaviour of mirror-image nucleic acid nanomaterials. First, L-DNA oligonucleotides can self-assemble into structurally defined nanoscale architectures with colloidal properties, similarly to native D-DNA nanostructures. Second, the sensitivity of the nanocube to dehydration and mechanical perturbation highlights the importance of preserving gentle aqueous conditions during handling and processing of highly ordered nucleic acid assemblies. Finally, from a translational perspective, these results also suggest that mirror-image nucleic acid nanostructures may require formulation strategies that minimize solvent-induced collapse or mechanical disruption during large-scale preparation and storage.

### Targeting and internalization of folic acid-functionalized L-DNA nanocube in cancer cells

Efficient intracellular delivery is a critical requirement for therapeutic nucleic acid nanostructures. To enable selective targeting of cancer cells, the mirror-image L-DNA nanocube was functionalized with folic acid (FA) through a click-compatible conjugation strategy (Fig. 2A and 2B). Folic acid was selected as a targeting ligand because folate receptors are overexpressed in a broad range of human cancers while remaining relatively low in many non-malignant cells^36^. By incorporating FA onto the surface of the biologically persistent L-DNA nanocube, we sought to combine receptor-mediated cellular uptake with the enhanced stability of mirror-image nucleic acid architectures. To generate FA-functionalized nanocubes, folic acid derivatives containing azide-reactive groups were conjugated onto pre-assembled L-DNA nanocubes bearing complementary click handles (Fig. 2C). This modular design enabled programmable surface functionalization without disrupting nanocube assembly or structural integrity. The resulting FA-L-DNA nanocubes were subsequently labelled with fluorescent probes to evaluate cellular binding and internalization.

**Figure 2.**
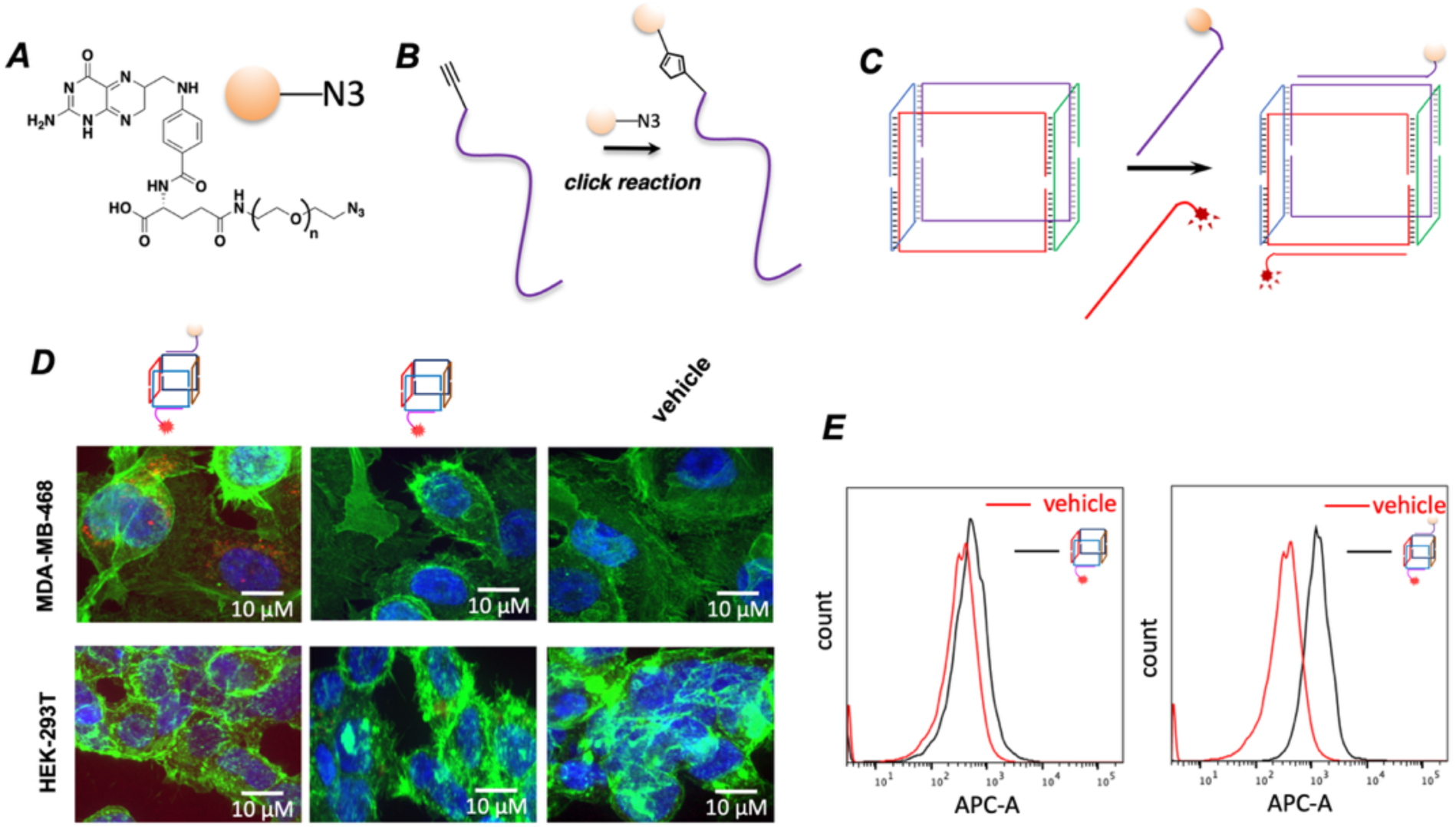
Binding and Internalization of L-DNA nanostructure mediated by the FA group. (A) Chemical structure of FA-(PEG)_n_-azido compound used for labelling L-DNA. (B) Click reaction to label 5’-alkyne modified L-DNA. (C) Schematic presentation of construction of the Cy5^+^, FA^+^ L-nanocube, containing FA (orange ball) targeting FA receptor and Cy5 fluorophore (red). (D) Confocal microscope images indicating the binding and internalization of L-nanocube into MDA-MB-468 cells (blue: nuclei; green: cytoskeleton; red: L-DNA nanostructures, the concentration of each fluorescent RNA nanostructures in treatment is 50 nM). (E) Flow cytometry analysis of Cy5-positive cells after treatment with Cy5⁺, FA⁺ L-nanocube and Cy5⁺, FA⁻ L-nanocube. The presence of folic acid significantly increased the efficiency of nanostructure’s binding to cancer cells.

The targeting specificity of FA-functionalized nanocubes was investigated using MDA-MB-468 breast cancer cells and HEK-293T cells as a comparatively low-folate-receptor control cell line. Media containing 50 nM Cy5 and FA labelled nanostructure was used to treat the cell lines. For comparison, the same incubations were performed using the Cy5 labelled nanostructures without the FA labelling, to validate the specific targeting. The cell culture media for drug treatment and cell growth contained Dulbecco’s Modified Eagle Medium and Fetal Bovine Serum without heat inactivation, to test the efficiency of L-DNA. Following the nanostructure incubation for 2h and washing steps, the cells were fixed, the nuclei were stained with DAPI, and the cytoskeletons were stained with FITC-conjugated phalloidin. Confocal fluorescence imaging revealed substantially enhanced uptake of FA-L-DNA nanocubes in MDA-MB-468 cells relative to HEK-293T cells (Fig. 2D). Strong intracellular fluorescence signals were observed throughout the cytoplasmic region of MDA-MB-468 cells following treatment with FA-functionalized nanocubes, whereas HEK-293T cells exhibited only weak fluorescence under identical conditions. As the comparison, the Cy5-positive, FA-negative L-DNA were not binding or delivered into either cancer cells or normal cells, due to the lack of FA molecule to direct cell targeting. Vehicle-treated control cells showed minimal background fluorescence in both cell lines.

Flow cytometric analysis further supported the confocal imaging results. MDA-MB-468 cells treated with Cy5-positive, FA-positive L-DNA nanocubes exhibited a pronounced rightward shift in APC fluorescence intensity relative to control groups, indicating efficient nanocube uptake (Fig. 2E, 68.2% of cells were Cy5-positive). In contrast, MDA-MB-468 cells treated with Cy5-positive, FA-negative L-DNA nanocubes displayed substantially lower fluorescence signals following treatment (22.7% of cells were Cy5-positive). These findings are consistent with folate receptor-mediated internalization and demonstrate that surface functionalization with FA significantly enhances selective cellular uptake of mirror-image nanocubes in cancer cells.

Importantly, the ability to chemically install targeting ligands onto the L-DNA scaffold highlights the modularity of the mirror-image nanocube platform. These results establish that mirror-image L-DNA nanostructures can be programmably functionalized with tumour-targeting ligands while preserving their structural integrity and biological stability. More broadly, the combination of programmable surface chemistry and intrinsic nuclease resistance supports the use of L-DNA nanocubes as versatile intracellular delivery platforms for targeted therapeutic applications.

### In vivo fluorescence imaging reveals enhanced tumour accumulation and persistence of L-DNA nanocubes

To evaluate the in vivo targeting capability and biological persistence of the mirror-image nanocube platform, tumour-bearing mice were intravenously administered folic acid and fluorescently labelled L-DNA nanocubes, D-DNA nanocube control, and PBS buffer, followed by longitudinal whole-body fluorescence imaging (Fig. 3). Mice received three systemic injections over a 6-day period, and fluorescence signals were monitored at multiple time points following the final administration.

**Figure 3.**
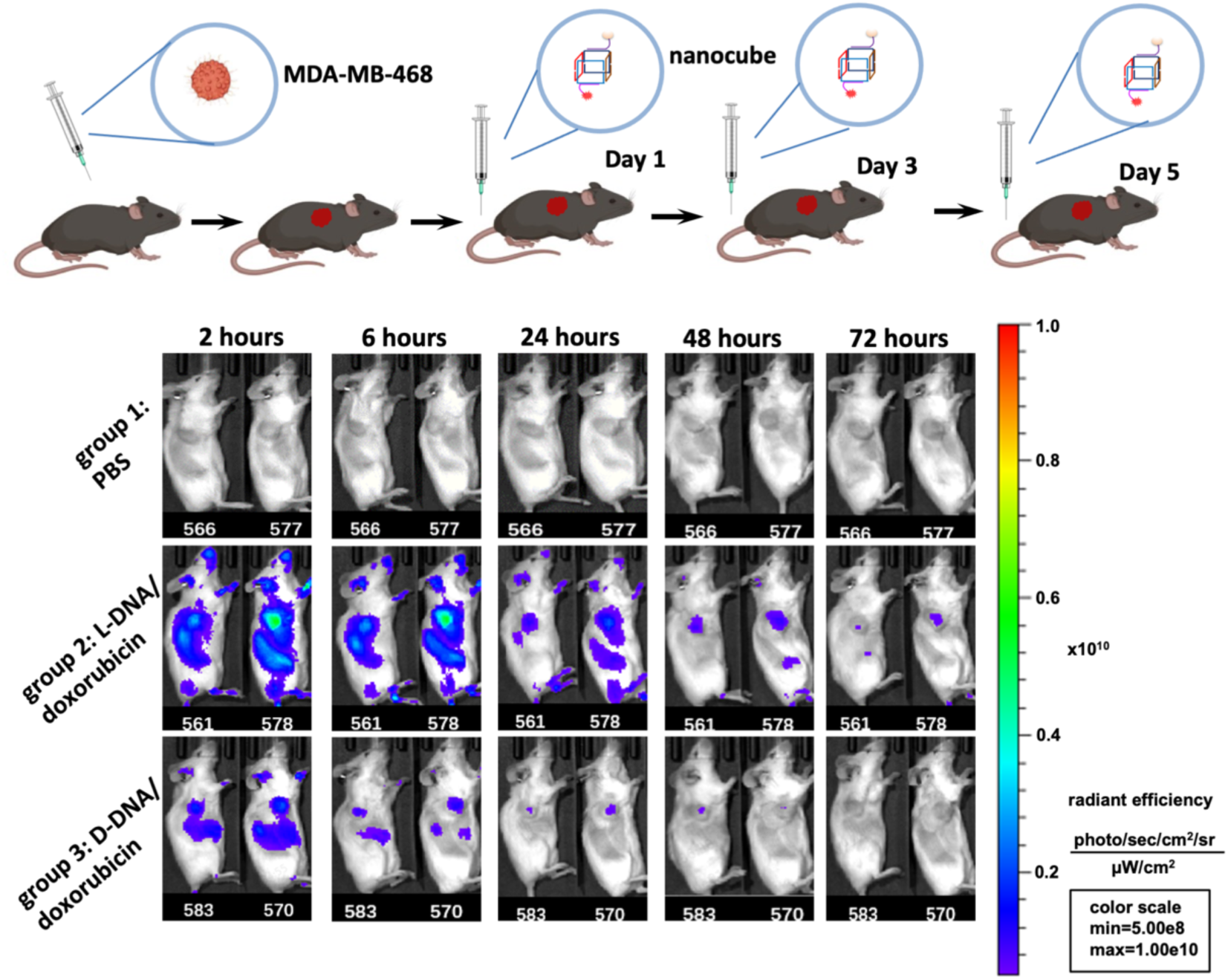
In vivo tumour targeting of mirror-image L-DNA nanocubes. (A) Schematic of systemic administration and fluorescence imaging of tumour-bearing mice treated with fluorescently labelled nanocubes. (B) Representative in vivo fluorescence images of mice treated with PBS, D-DNA nanocubes or L-DNA nanocubes at the indicated time points.

Consistent with the enhanced biological stability of the mirror-image scaffold, L-DNA nanocubes exhibited substantially prolonged in vivo fluorescence retention compared with corresponding D-DNA nanocubes. Strong fluorescence signals were detected in tumour regions of mice treated with L-DNA nanocubes over multiple imaging time points, whereas fluorescence associated with D-DNA nanocubes decreased more rapidly following administration. The persistent fluorescence observed in tumour tissue suggests that the mirror-image nanocube remains structurally intact during systemic circulation and retains the ability to undergo receptor-mediated tumour targeting following intravenous administration. In contrast, the reduced signal intensity observed for D-DNA nanocubes is consistent with partial degradation and destabilization of the native nanostructure under physiological conditions.

The improved in vivo stability of the L-DNA nanocube likely arises from multiple cooperative effects associated with mirror-image chirality. First, resistance to endogenous serum nucleases enables the nanocube to remain structurally intact during circulation, thereby preserving the integrity of the targeted delivery scaffold. Second, prolonged biological persistence increases the probability of tumour accumulation through both receptor-mediated uptake and extended systemic exposure. These in vivo imaging results further establish that mirror-image L-DNA nanostructures can function as stable and programmable delivery scaffolds in living systems. The combination of tumour-selective targeting, prolonged circulation-associated persistence and enhanced in vivo stability supports the use of L-DNA nanocubes as versatile therapeutic nanomaterials for intracellular drug delivery applications.

### Doxorubicin-loaded L-DNA nanocubes inhibit cancer cell growth

To investigate the therapeutic potential of the mirror-image nanocube platform, doxorubicin (DOX) was incorporated into the L-DNA nanocube through intercalation-based loading (Fig. 4). Because anthracycline drugs such as DOX^37, 38^.exhibit strong interactions with nucleic acid duplex regions, the L-DNA nanocube provides a structurally defined scaffold capable of carrying multiple DOX molecules while maintaining nanoscale organization and biological stability (Fig. 4A). We hypothesized that the enhanced persistence and tumour-targeting capability of the mirror-image nanocube would improve intracellular delivery of DOX while reducing nonspecific toxicity.

**Figure 4.**
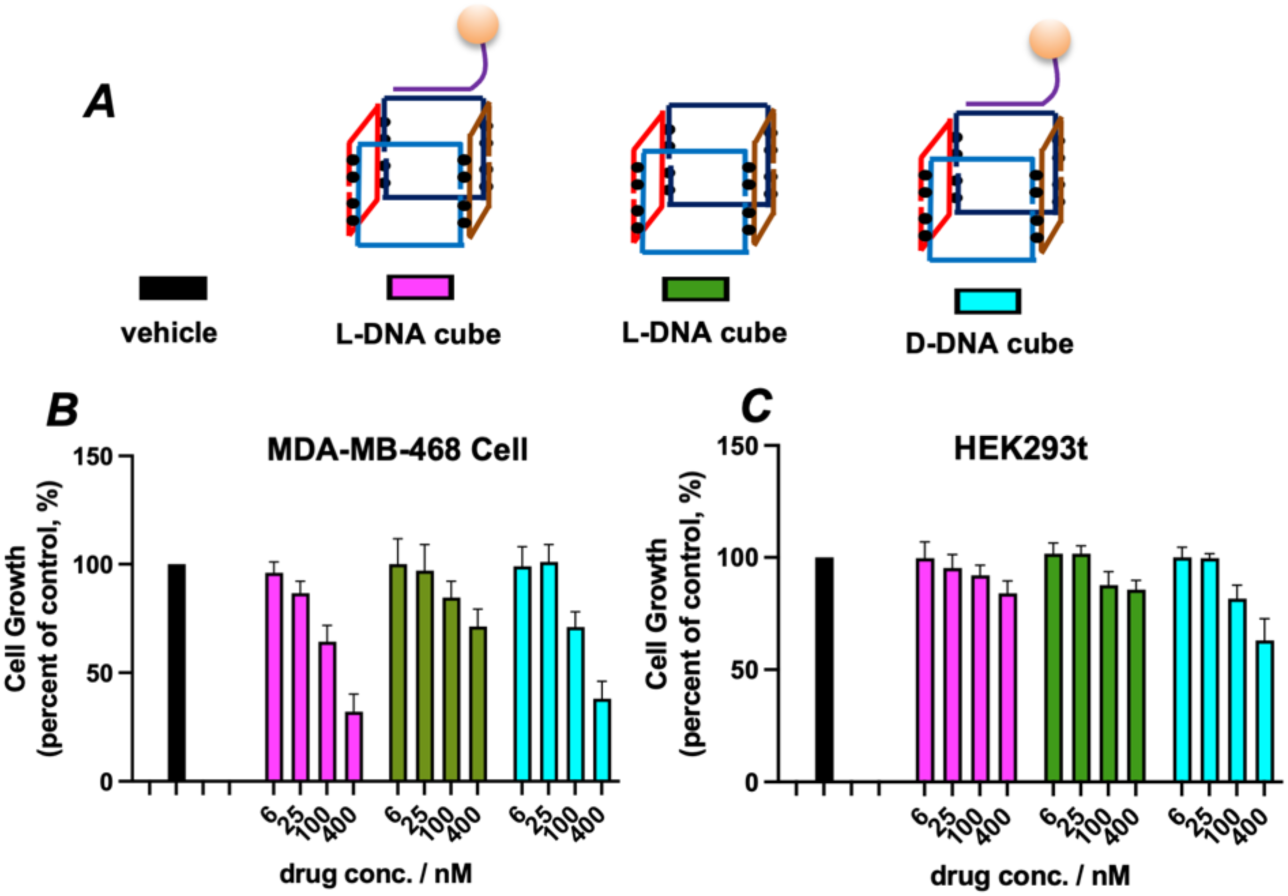
Comparison of inhibition of cell growth in MDA-MB-468 and HEK293t cells by three nanocubes. (A) Schematic representations of L-DNA nanocubes with and without FA conjugation, and D-DNA nanocube with FA conjugation. Black dots: DOX; orange ball: FA group. (B) Alamar blue assay using different nanocubes to treat MDA-MB-468 at 4 different concentrations. (C) Alamar blue assay using different nanocubes to treat HEK293t cells at 4 different concentrations. Data are representative of triplicate experiments. Error bars represent the mean ± SEM from three independent experiments (n = 3).

The cytotoxic effects of DOX-loaded nanocubes were evaluated in cancer cells using Alamar Blue viability assays. For the viability assay of each drug, we performed the experiment at 4 different concentrations, subjected to serial dilutions by a factor of four. The y-axis indicates the relative cell viability compared to the control group, where no drug was added. Cells treated with FA and DOX-loaded L-DNA nanocubes exhibited substantially reduced viability relative to vehicle-treated controls and empty nanocube formulations, indicating efficient intracellular delivery and pharmacological activity of the loaded drug (Fig. 4B). Folic acid enhances receptor-mediated uptake in cancer cells, while doxorubicin contributes to cytotoxicity through DNA intercalation and inhibition of topoisomerase II. We observe these nanostructures suppressed cancer cell growth by ∼70% at the 400 nM concentration treatment. As comparison, the DOX-loaded nanocube without FA presents reduced inhibitory ability, which indicates the function of FA in cancer cell targeting.

We then performed the same experiment in HEK293t cells, which do not express the folic acid receptor on their cell membranes. L-DNA nanostructures with or without folic acid did not exhibit strong toxicity, especially at low concentration. This is because of the absence of FA-directed drug delivery due to the lack of FA-receptor interaction. Only weak toxicity was observed at 400 nM, inhibiting cell proliferation by approximately 20%. Considering the fact that HEK293t cells are lack of FA receptors on the cell membrane to facilitate efficient endocytosis, the observed toxicity is likely due to the doxorubicin leaking into the media during incubation, followed by passive penetration across the cell membrane. Notably, FA- and DOX-loaded D-DNA nanocubes exhibited greater cytotoxicity than the corresponding L-DNA nanocubes at the highest tested concentration (400 nM). This difference is likely attributable to the lower biological stability of the D-DNA scaffold under cell culture conditions, which may promote partial nanocube dissociation and premature release of free DOX into the extracellular environment. In contrast, the enhanced structural persistence of the mirror-image L-DNA nanocube likely limits uncontrolled drug leakage, thereby reducing nonspecific toxicity while maintaining intracellular delivery capability. These observations further support the improved biological stability and potential biosafety advantages of the L-DNA nanocube platform.

Overall, our findings further demonstrate that the L-DNA nanocube can function as a multifunctional therapeutic scaffold capable of incorporating small-molecule drugs while preserving structural integrity and intracellular delivery capability. More broadly, the results establish that mirror-image nucleic acid nanostructures can support biologically active chemotherapeutic delivery without requiring conventional lipid or polymer carrier systems.

### L-DNA nanocube-mediated delivery reduces systemic toxicity associated with free doxorubicin treatment

To assess the in vivo safety profile of the L-DNA nanocube drug delivery platform, tumour-bearing mice were treated with PBS, L-DNA/DOX nanocubes, D-DNA/DOX nanocubes or free doxorubicin (DOX), followed by evaluation of plasma urea levels and histological examination of major organs (Fig. 5A). Because DOX is known to cause dose-limiting systemic toxicity, particularly in highly perfused organs such as the kidney, these analyses were used to compare the relative safety profiles of free drug and nanocube-associated delivery.

**Figure 5.**
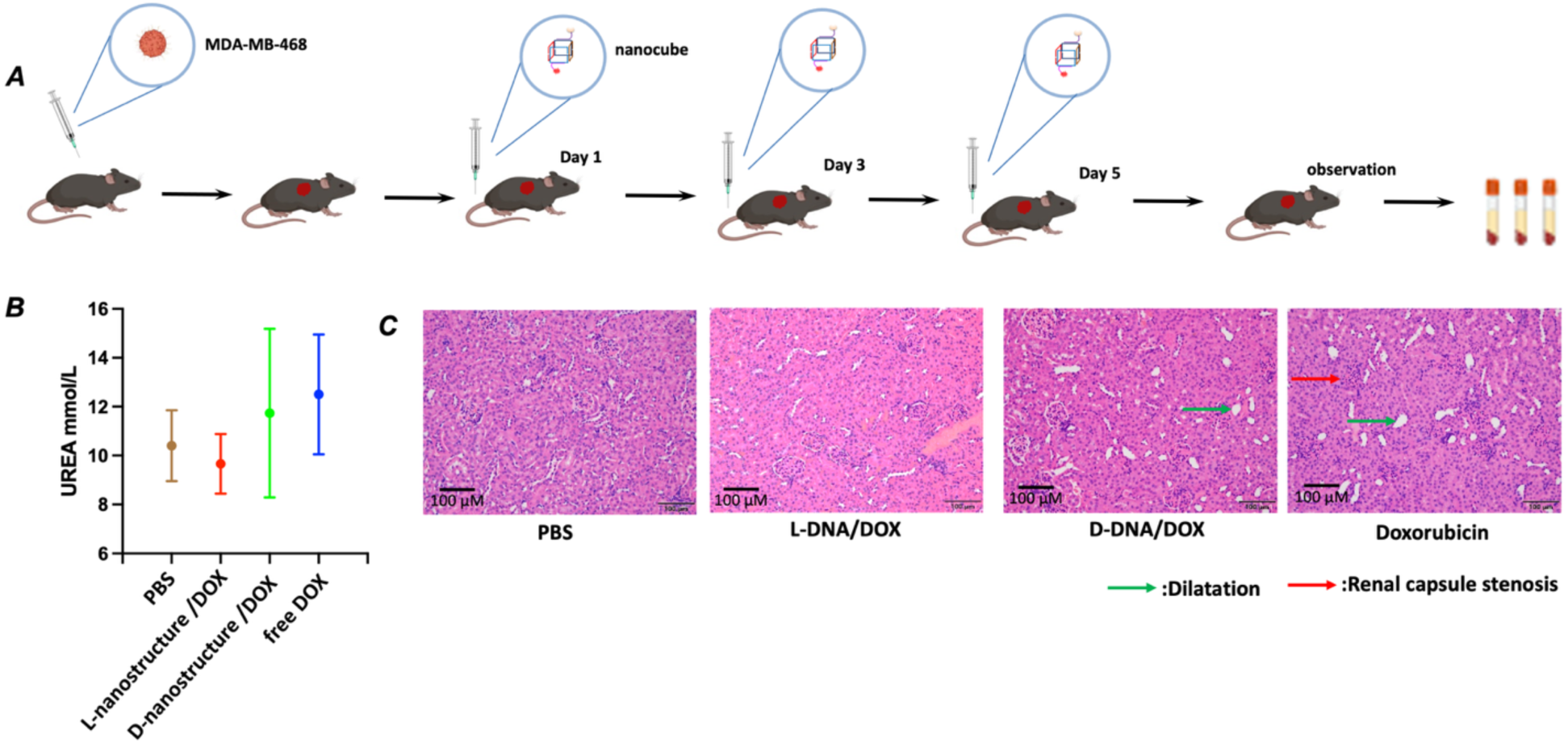
In vivo toxicity evaluation of doxorubicin-loaded mirror-image nanocubes. (A) Experimental design for systemic administration of PBS, free DOX, D-DNA/DOX nanocubes and L-DNA/DOX nanocubes in tumour-bearing mice. (B) Plasma urea measurements following treatment with the indicated formulations. Data are presented as mean ± s.d. (C) Representative H&E-stained kidney sections from mice treated with PBS, free DOX, D-DNA/DOX nanocubes or L-DNA/DOX nanocubes. Histological alterations were more apparent in free DOX and D-DNA/DOX groups, whereas kidney morphology in the L-DNA/DOX group appeared comparatively preserved.

Plasma urea measurements showed only modest differences among the treatment groups, with no dramatic elevation detected in the L-DNA/DOX group relative to PBS controls. The D-DNA/DOX and free DOX groups showed a trend toward higher or more variable urea values, although the differences remained limited and did not support a strong conclusion of overt renal protection (Fig. 5B). These data suggest that nanocube-mediated delivery did not increase systemic renal burden and may have helped moderate acute renal stress compared with free DOX exposure.

Histological analysis of kidney sections by H&E staining provided additional context for the biochemical measurements (Fig. 5C). PBS-treated animals showed preserved renal architecture, whereas free DOX-treated mice displayed more apparent histological abnormalities, including renal capsule stenosis and structural distortion. The D-DNA/DOX group also showed visible renal changes, including tubular or tissue dilation, whereas the L-DNA/DOX group appeared comparatively closer to the PBS control, with fewer obvious histopathological alterations. These observations are consistent with, but do not definitively prove, a reduced renal toxicity profile for L-DNA-mediated DOX delivery.

The modest differences across groups is not unexpected, given the small sample size and the fact that plasma urea and routine H&E are relatively coarse readouts of toxicity. Nevertheless, the combined biochemical and histological trends support the view that the mirror-image nanocube platform may alter the systemic exposure profile of doxorubicin in a way that is less damaging to renal tissue than free drug administration. More importantly, these data suggest that the enhanced biological persistence of the L-DNA scaffold does not appear to worsen acute organ toxicity under the conditions tested. Overall, this toxicity study should be interpreted as supportive rather than definitive. The results are consistent with a potentially improved safety profile for L-DNA nanocube-mediated DOX delivery, but additional studies incorporating larger cohorts, expanded serum chemistry, and longer-term histopathology will be needed to fully define whether the mirror-image nanocube platform provides a reproducible toxicity advantage over free doxorubicin or D-DNA controls.

### pH-responsive intracellular release promotes cytosolic siRNA delivery

To facilitate intracellular release of therapeutic cargo, we engineered the L-DNA nanocube with a folic acid-conjugated L-DNA, and a pH-responsive sequence capable of forming a triplex structure under acidic conditions (Figure 6A). Because endosomal maturation is accompanied by progressive acidification, we hypothesized that triplex formation within the endosomal environment would destabilize siRNA hybridization and trigger cargo release from the nanocube scaffold. In addition, the released siRNA complex contains a membrane-active peptide designed to promote endosomal escape following intracellular dissociation. The peptide we choose to conjugate with siRNA is the pH-responsive fusogenic peptide INF7^39^, a derivative of the influenza virus hemagglutinin HA2 fusion peptide that undergoes conformational activation under acidic conditions and promotes endosomal membrane destabilization^40^. The combination of triplex-mediated siRNA release and INF7-assisted membrane disruption was designed to facilitate cytosolic delivery following endosomal acidification.

**Figure 6.**
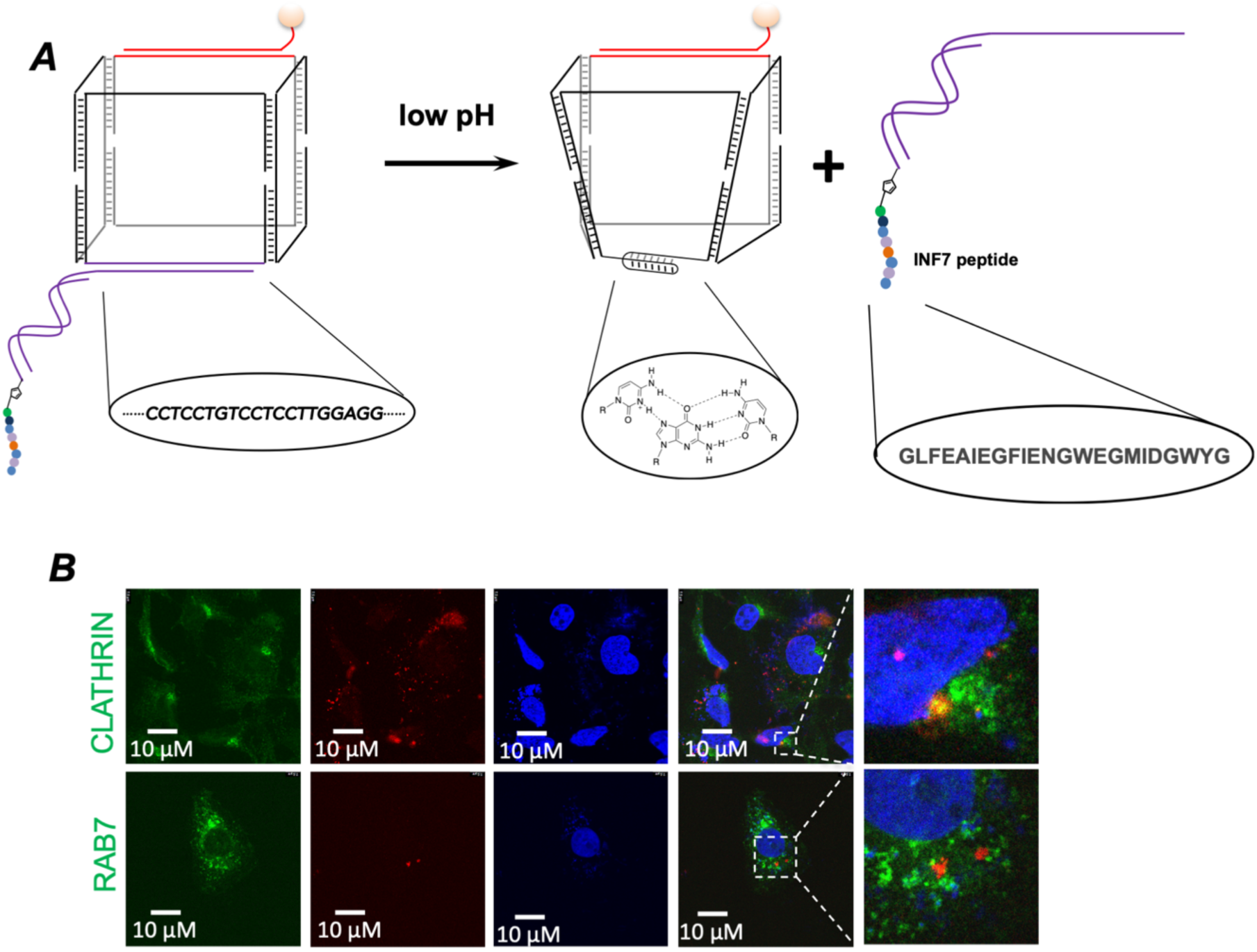
pH-responsive intracellular release and trafficking of siRNA delivered by L-DNA nanocubes. (A) Schematic illustration of the pH-responsive siRNA delivery mechanism. Following folate receptor-mediated endocytosis, endosomal acidification induces triplex formation within the nanocube scaffold, resulting in release of the siRNA-INF7 conjugate and subsequent intracellular redistribution. (B) Confocal fluorescence images of cells treated with siRNA-loaded L-DNA nanocubes and co-stained with clathrin or the late endosomal marker RAB7. Released siRNA exhibited limited colocalization with endocytic compartments following intracellular delivery. Nuclei were stained with DAPI.

To investigate intracellular trafficking following nanocube uptake, cancer cells were incubated with fluorescently labelled siRNA-loaded L-DNA nanocubes and analysed by confocal microscopy. The nanocube surface was functionalized with folic acid to promote receptor-mediated endocytosis through folate receptor-positive cancer cells, thereby facilitating intracellular accumulation of the therapeutic construct. Following endocytic internalization, progressive acidification within endosomal compartments was expected to induce triplex formation by the engineered pH-responsive sequence embedded within the nanocube scaffold. This structural rearrangement was designed to destabilize siRNA hybridization and trigger release of the siRNA-INF7 conjugate from the nanocube architecture. To evaluate the intracellular trafficking of released siRNA, cells were co-stained with markers associated with distinct stages of endocytic trafficking, including clathrin^41^ and the late endosomal marker RAB7^42^. Confocal imaging revealed minimal colocalization between siRNA signals and either clathrin- or RAB7-positive compartments following intracellular delivery (Fig. 6B). The limited overlap with clathrin or RAB7 suggests that a substantial fraction of released siRNA was not retained within early or late endosomal structures. These observations support the proposed mechanism in which endosomal acidification triggers intracellular cargo dissociation, followed by redistribution of released siRNA away from canonical endocytic compartments. Our observed trafficking behavior is consistent with the functional role of INF7, a pH-responsive fusogenic peptide derived from the influenza hemagglutinin HA2 domain. Under acidic endosomal conditions, INF7 undergoes conformational activation that enhances membrane destabilization and facilitates translocation of cargo across endosomal membranes. The combination of triplex-mediated siRNA release and INF7-assisted membrane disruption therefore provides a sequential intracellular activation mechanism that couples environmental sensing with cytosolic delivery.

### Immunogold TEM analysis supports intracellular redistribution of delivered siRNA

To further investigate the intracellular fate of siRNA following L-DNA nanocube-mediated delivery, we performed immunogold transmission electron microscopy (TEM) using biotinylated siRNA detected with streptavidin-conjugated nanogold particles (Fig. 7A). The nanocube platform was engineered to promote sequential intracellular activation following receptor-mediated endocytosis. Specifically, folic acid functionalization facilitated tumour cell uptake, whereas progressive acidification within endosomal compartments was designed to induce triplex formation by the pH-responsive nanocube sequence, thereby triggering release of the siRNA-INF7 conjugate. The released INF7-modified siRNA was expected to subsequently promote membrane destabilization and intracellular redistribution through the pH-dependent fusogenic activity of INF7.

**Figure 7.**
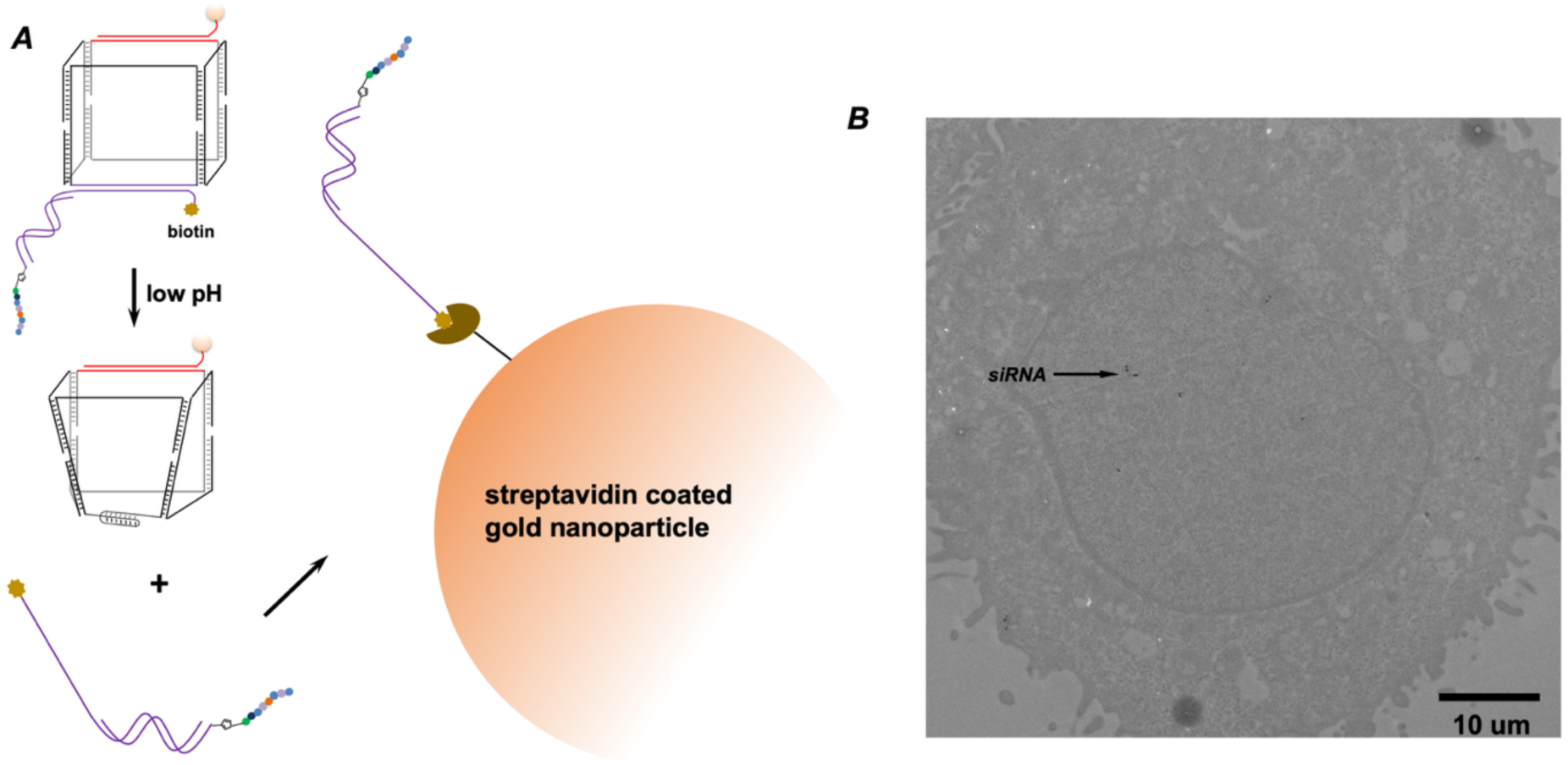
Immunogold TEM analysis of intracellular siRNA localization following L-DNA nanocube delivery.. (A) Schematic illustration of biotinylated siRNA release from the L-DNA nanocube following endosomal acidification and subsequent detection using streptavidin-conjugated nanogold particles for TEM visualization. (B) Representative immunogold TEM image of cells treated with siRNA-loaded L-DNA nanocubes. Electron-dense nanogold puncta corresponding to delivered biotinylated siRNA were detected within cytoplasmic regions Control samples showed no detectable signal (supporting information).

Initial immunogold labeling strategies produced substantial nonspecific background signals distributed across multiple intracellular compartments, limiting reliable interpretation of intracellular siRNA localization. To improve labeling specificity, we optimized the electron microscopy workflow using post-embedding immunogold labeling conditions without osmium tetroxide (OsO4) treatment, combined with biotinylated siRNA detection through streptavidin-conjugated nanogold particles. Under these optimized conditions, electron-dense nanogold puncta were selectively detected in siRNA-treated samples, whereas control cells processed in parallel showed no detectable signal (Fig. 7B). Although the overall labeling density remained relatively sparse, which is likely attributable to the limited abundance of intracellularly released siRNA together with incomplete labeling efficiency during post-embedding detection, discrete nanogold puncta were observed within cytoplasmic regions outside clearly defined membrane-bound vesicular structures. The low background observed in control samples supports the specificity of the optimized labeling strategy. These observations are consistent with intracellular redistribution of released siRNA following nanocube uptake and endosomal acidification.

The relatively sparse distribution of immunogold labeling is also biologically plausible given the transient and heterogeneous nature of endosomal escape during intracellular nucleic acid delivery. Efficient cytosolic release of siRNA is generally considered a low-frequency event for many nanoparticle systems, and direct ultrastructural visualization of released intracellular nucleic acid cargo remains technically challenging. In this context, the detection of treatment-specific cytoplasmic nanogold puncta, together with the reduced colocalization of siRNA with RAB7-positive endosomal compartments observed by confocal microscopy and the subsequent gene silencing activity, provides convergent evidence supporting functional intracellular delivery by the L-DNA nanocube platform. These findings further support the proposed mechanism in which the mirror-image nanocube architecture functions not only as a biologically persistent delivery scaffold, but also as a programmable intracellular release system capable of coupling environmental pH sensing with membrane-active cargo redistribution.

### Chemically programmable bortezomib prodrug conjugation enables intracellular delivery of bortezomib

To further expand the chemical scope of the mirror-image L-DNA nanocube platform, we designed three structurally distinct bortezomib (BTZ) prodrugs based on the boronic acid pharmacophore of BTZ. In each case, the reactive boronic acid motif was masked as a boronate ester and functionalized with an azide handle to enable post-assembly click conjugation to alkyne-modified L-DNA nanocubes (Fig. 8A-8C). This design was intended to improve the stability of the native BTZ pharmacophore during extracellular exposure and nanoparticle handling, while preserving the possibility of intracellular reactivation after cellular uptake (Fig. 8D). After nanocube assembly, the three azide-functionalized BTZ prodrugs were covalently installed onto pre-formed L-DNA nanocubes through copper-catalysed azide–alkyne cycloaddition, and excess small-molecule species were removed by size exclusion chromatography.

**Figure 8.**
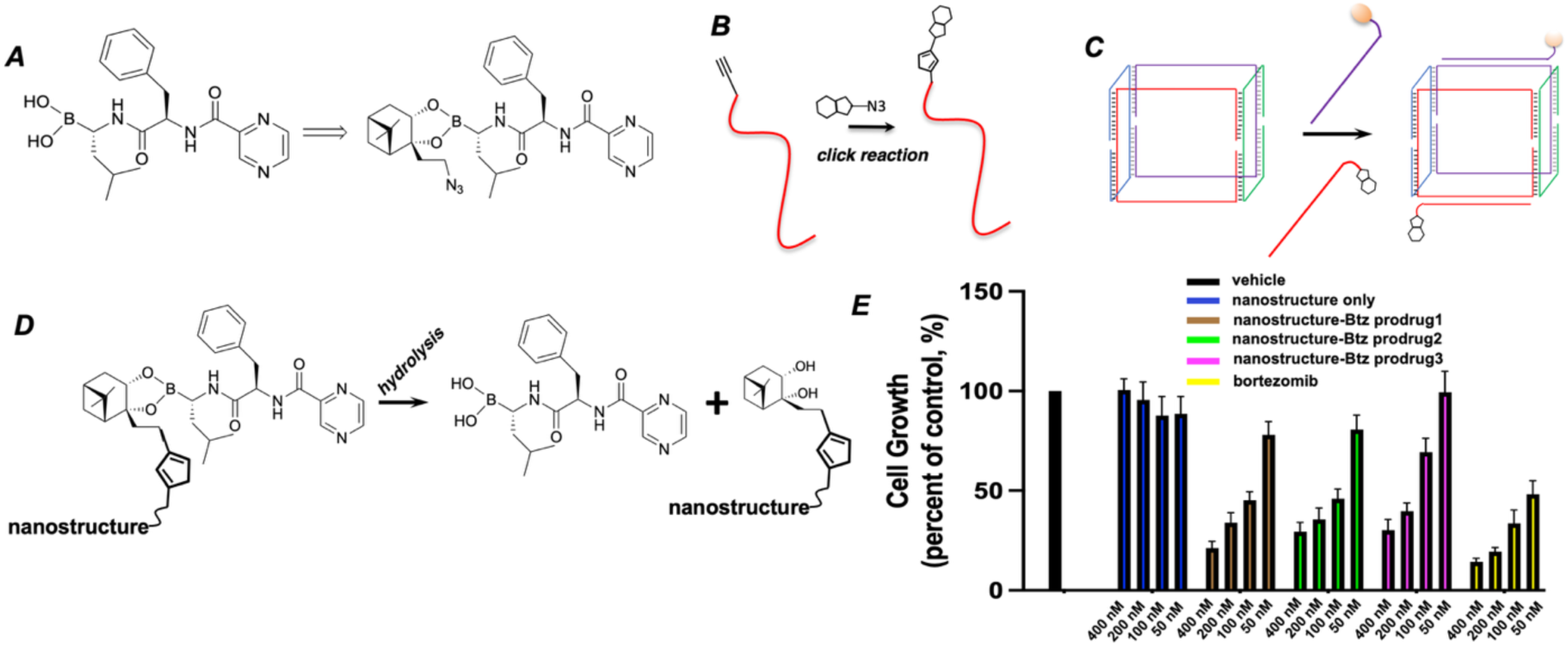
Covalent conjugation and intracellular delivery of bortezomib prodrugs using L-DNA nanocubes. (A) Chemical structures of bortezomib and one of the three azide-functionalized bortezomib boronate ester prodrugs used in this study. (B) Schematic illustration of azide-functionalized bortezomib boronate ester prodrugs conjugated onto alkyne-functionalized L-DNA through copper-catalysed azide–alkyne cycloaddition (CuAAC) chemistry. (C) Schematic representation of functionalization of L-DNA nanocube with bortezomib prodrugs and folic acid. (D) Proposed intracellular activation mechanism of boronate ester hydrolysis and release of active bortezomib following nanocube-mediated delivery. (E) Alamar Blue cell viability analysis of cancer cells treated with the indicated bortezomib prodrug-functionalized L-DNA nanocubes, free bortezomib or nanocube-only controls. Data are presented as mean ± s.d. from three independent experiments (n = 3).

All three BTZ prodrug candidates exhibited measurable cytotoxicity in Alamar Blue assays after nanocube-mediated delivery, indicating that the conjugated prodrug architectures remained biologically active and could be converted into pharmacologically relevant species intracellularly (Fig. 8E). At the same time, the toxicity of the prodrug-loaded nanocubes was consistently lower than that of free BTZ at equivalent nominal concentrations. This difference is expected, because the boronate ester prodrug architecture introduces an additional activation barrier relative to free BTZ: the boronic acid is transiently masked, intracellular hydrolysis is required to regenerate active drug, and nanocube trafficking adds an extra delivery step before target engagement. In addition, the three prodrugs were designed with distinct boronate ester structures, which likely differ in hydrolytic stability, steric accessibility and release kinetics. A more stable boronate ester is expected to improve systemic handling and premature-drug suppression, whereas a less stable ester may release BTZ more rapidly once internalized. The observed intermediate toxicity profile is therefore consistent with a tunable prodrug platform in which chemical stability and intracellular release can be modulated by linker architecture.

Mechanistically, the lower toxicity relative to free BTZ is also reasonable from a delivery perspective. Free BTZ can rapidly access intracellular targets without requiring endocytic uptake or prodrug hydrolysis, whereas the nanocube-conjugated prodrug must first undergo receptor-mediated internalization, endosomal trafficking and hydrolytic unmasking before active BTZ becomes available. As a result, the nanocube formulation is expected to attenuate immediate pharmacological potency while improving formulation stability and enabling more controlled intracellular drug activation. This trade-off is advantageous for therapeutic delivery, because it may reduce off-target toxicity associated with premature exposure of the active boronic acid pharmacophore while still retaining sufficient intracellular activity to inhibit cancer cell growth.

Importantly, this BTZ study extends the utility of the L-DNA nanocube beyond simple cargo loading and demonstrates that mirror-image nucleic acid nanostructures can serve as a chemically programmable scaffold for covalent delivery of boronic acid-containing therapeutics. More broadly, these results provide a generalizable strategy for the delivery of boronic acid-based drugs, a class of molecules whose pharmacological performance is often limited by reactivity, hydrolytic instability and premature target engagement. By combining boronate ester masking, click-conjugation chemistry and a biologically persistent L-DNA carrier, this platform offers a modular route to intracellular delivery of boronic acid therapeutics with tunable stability and activation profiles.

## CONCLUSION

In summary, we report a mirror-image L-DNA nanocube as a biologically persistent and chemically programmable platform for intracellular therapeutic delivery. The self-assembled L-DNA nanostructure preserves the structural programmability of conventional nucleic acid nanotechnology while exhibiting substantially enhanced resistance to enzymatic degradation and improved stability under biologically relevant conditions. Through modular surface engineering, the nanocube can be functionalized with targeting ligands such as folic acid to promote selective tumor cell uptake and enhanced in vivo tumor accumulation. Beyond structural stability, the L-DNA nanocube supports delivery of chemically diverse therapeutic cargos, including intercalated chemotherapeutic agents, covalently conjugated boronic acid-based prodrugs and functional siRNA payloads. The platform further enables programmable intracellular activation through pH-responsive structural rearrangement and INF7-assisted intracellular redistribution, facilitating cytosolic delivery of siRNA and subsequent gene silencing activity. Importantly, the mirror-image scaffold maintained therapeutic functionality while exhibiting trends toward reduced nonspecific toxicity relative to conventional D-DNA nanostructures and free drug controls.

Our findings establish that mirror-image nucleic acid nanostructures can function not only as stable delivery carriers, but also as biologically functional nanomaterials with programmable chemical and intracellular behaviors. More broadly, this work introduces a general strategy for integrating biological persistence, modular cargo engineering and environmentally responsive intracellular activation within a single nucleic acid nanotechnology platform. We anticipate that mirror-image nucleic acid nanostructures may provide new opportunities for the development of multifunctional therapeutic systems and expand the translational scope of structural nucleic acid nanotechnology.

## SUPPORTING INFORMATION

Additional experimental details, materials and methods, including sequences of the synthetic RNAs and their MS characterizations.

## ACKNOWLEDGEMENTS

We thank Dr. Zhang and members for helpful discussions, insightful commentary, and careful revision of the manuscript. We thank Dr. L. Zeng and the Chemical Genomics Core at IUSM for spectropolarimeter analysis. We thank Dr. Giovanni Gonzalez-Gutierrez in the Physical Biochemistry Instrumentation Core, IU Bloomington for the DLS study. We thank Drs. Anthony Sinn and Felicia Kennedy from Laboratory Animal Resource Center at Indiana University School of Medicine for the animal study. We thank Drs. Quyen Quoc Hoang and Yanshin Park from Center for Electron Microscopy (iCEM) at Indiana University School of Medicine for the TEM study. This work was supported in part by 100 Voices of Hope award from the Indiana University to W.Z., and by Showalter Trust Award from Showalter Trust Foundation to W.Z.

## CONFLICT OF INTEREST

The authors declare no conflicts of interest.

## Notes

### Competing Interest Statement

The authors have declared no competing interest.

## REFERENCES

(1) Guo, P. The emerging field of RNA nanotechnology. Nat. Nanotechnol. 2010, 5 (12), 833–842.

(2) Jasinski, D.; Haque, F.; Binzel, D. W.; Guo, P. Advancement of the emerging field of RNA nanotechnology. ACS Nano 2017, 11 (2), 1142–1164.

(3) Guo, P. RNA nanotechnology: engineering, assembly and applications in detection, gene delivery and therapy. J. Nanosci. Nanotechnol. 2005, 5 (12), 1964–1982.

(4) Krishnan, Y.; Seeman, N. C. Introduction: nucleic acid nanotechnology. ACS Publications: 2019; Vol. 119, pp 6271–6272.

(5) Tsui, N. B.; Ng, E. K.; Lo, Y. D. Stability of endogenous and added RNA in blood specimens, serum, and plasma. Clin. Chem. 2002, 48 (10), 1647–1653.

(6) Abdelmawla, S.; Guo, S.; Zhang, L.; Pulukuri, S. M.; Patankar, P.; Conley, P.; Trebley, J.; Guo, P.; Li, Q.-X. Pharmacological characterization of chemically synthesized monomeric phi29 pRNA nanoparticles for systemic delivery. Mol. Ther. 2011, 19 (7), 1312–1322.

(7) Paredes, E.; Evans, M.; Das, S. R. RNA labeling, conjugation and ligation. Methods 2011, 54 (2), 251–259.

(8) Xie, X.; Ma, W.; Zhan, Y.; Zhang, Q.; Wang, C.; Zhu, H. Methods to improve the stability of nucleic acid-based nanomaterials. Current Drug Metabolism 2023, 24 (5), 315–326.

(9) Burnett, J. C.; Rossi, J. J. RNA-based therapeutics: current progress and future prospects. Chem. Biol. 2012, 19 (1), 60–71.

(10) Singha, K.; Namgung, R.; Kim, W. J. Polymers in small-interfering RNA delivery. Nucleic acid therapeutics 2011, 21 (3), 133–147.

(11) Brader, M. L.; Williams, S. J.; Banks, J. M.; Hui, W. H.; Zhou, Z. H.; Jin, L. Encapsulation state of messenger RNA inside lipid nanoparticles. Biophysical journal 2021, 120 (14), 2766–2770.

(12) Sokolova, V.; Epple, M. Inorganic nanoparticles as carriers of nucleic acids into cells. Angewandte chemie international edition 2008, 47 (8), 1382–1395.

(13) Shen, W.; De Hoyos, C. L.; Sun, H.; Vickers, T. A.; Liang, X.-h.; Crooke, S. T. Acute hepatotoxicity of 2′ fluoro-modified 5–10–5 gapmer phosphorothioate oligonucleotides in mice correlates with intracellular protein binding and the loss of DBHS proteins. Nucleic Acids Res. 2018, 46 (5), 2204–2217.

(14) Corey, D. R. Chemical modification: the key to clinical application of RNA interference? J. Clin. Invest. 2007, 117 (12), 3615–3622.

(15) Deleavey, G. F.; Damha, M. J. Designing chemically modified oligonucleotides for targeted gene silencing. Chemistry & Biology 2012, 19 (8), 937–954.

(16) Ashley, G. W. Modeling, synthesis, and hybridization properties of (L)-ribonucleic acid. J. Am. Chem. Soc. 1992, 114 (25), 9731–9736.

(17) Dantsu, Y.; Zhang, W. Derivatization of Mirror-image L-Nucleic Acids with 2’-OMe Modification for Thermal and Structural Stabilization. ChemBioChem 2023, e202200764.

(18) Dantsu, Y.; Zhang, Y.; Zhang, W. Synthesis of 2′-Deoxy-2′-fluoro-l-cytidine and Fluorinated l-Nucleic Acids for Structural Studies. ChemistrySelect 2021, 6 (39), 10597–10600.

(19) Dantsu, Y.; Zhang, Y.; Zhang, W. Synthesis and Structural Characterization of 2′-Deoxy-2′-fluoro-l-uridine Nucleic Acids. Org. Lett. 2021, 23 (13), 5007–5011.

(20) Kabza, A. M.; Sczepanski, J. T. An l-RNA Aptamer with Expanded Chemical Functionality that Inhibits MicroRNA Biogenesis. ChemBioChem 2017, 18 (18), 1824–1827.

(21) Young, B. E.; Kundu, N.; Sczepanski, J. T. Mirror-image oligonucleotides: history and emerging applications. Chem. Eur. J. 2019, 25 (34), 7981–7990.

(22) Dantsu, Y.; Zhang, Y.; Zhang, W. Advances in therapeutic L-nucleosides and L-nucleic acids with unusual handedness. Genes 2022, 13 (1), 46.

(23) Eulberg, D.; Klussmann, S. Spiegelmers: biostable aptamers. Chembiochem 2003, 4 (10), 979–983.

(24) Vater, A.; Klussmann, S. Turning mirror-image oligonucleotides into drugs: the evolution of Spiegelmer® therapeutics. Drug discovery today 2015, 20 (1), 147–155.

(25) Vater, A.; Klussmann, S. Turning mirror-image oligonucleotides into drugs: the evolution of Spiegelmer® therapeutics. Drug Discov. Today 2015, 20 (1), 147–155.

(26) Sczepanski, J. T.; Joyce, G. F. Specific inhibition of microRNA processing using L-RNA aptamers. J. Am. Chem. Soc. 2015, 137 (51), 16032–16037.

(27) Sczepanski, J. T.; Joyce, G. F. Binding of a structured D-RNA molecule by an L-RNA aptamer. J. Am. Chem. Soc. 2013, 135 (36), 13290–13293.

(28) Li, J.; Sczepanski, J. T. Targeting a conserved structural element from the SARS-CoV-2 genome using l-DNA aptamers. RSC Chem. Biol. 2022, 3 (1), 79–84.

(29) Thai, H. B. D.; Kim, K.-R.; Hong, K. T.; Voitsitskyi, T.; Lee, J.-S.; Mao, C.; Ahn, D.-R. Kidney-targeted cytosolic delivery of siRNA using a small-sized mirror DNA tetrahedron for enhanced potency. ACS Cent. Sci. 2020, 6 (12), 2250–2258.

(30) Sun, Y.; Yang, B.; Hua, Y.; Dong, Y.; Ye, J.; Wang, J.; Xu, L.; Liu, D. Construction and Characterization of a Mirror-Image l-DNA i-Motif. ChemBioChem 2020, 21 (1-2), 94–97.

(31) Yang, B.; Zhou, B.; Li, C.; Li, X.; Shi, Z.; Li, Y.; Zhu, C.; Li, X.; Hua, Y.; Pan, Y. A Biostable l-DNA Hydrogel with Improved Stability for Biomedical Applications. Angew. Chem. 2022, 134 (30), e202202520.

(32) Hueso, M.; Mallén, A.; Suñé-Pou, M.; Aran, J. M.; Suñé-Negre, J. M.; Navarro, E. ncRNAs in therapeutics: challenges and limitations in nucleic acid-based drug delivery. International journal of molecular sciences 2021, 22 (21), 11596.

(33) Dowdy, S. F.; Setten, R. L.; Cui, X.-S.; Jadhav, S. G. Delivery of RNA therapeutics: the great endosomal escape! Nucleic acid therapeutics 2022, 32 (5), 361–368.

(34) Rampersad, S. N. Multiple applications of Alamar Blue as an indicator of metabolic function and cellular health in cell viability bioassays. Sensors 2012, 12 (9), 12347–12360.

(35) Haldón, E.; Nicasio, M. C.; Pérez, P. J. Copper-catalysed azide–alkyne cycloadditions (CuAAC): an update. Organic & biomolecular chemistry 2015, 13 (37), 9528–9550.

(36) Cheung, A.; Bax, H. J.; Josephs, D. H.; Ilieva, K. M.; Pellizzari, G.; Opzoomer, J.; Bloomfield, J.; Fittall, M.; Grigoriadis, A.; Figini, M. Targeting folate receptor alpha for cancer treatment. Oncotarget 2016, 7 (32), 52553.

(37) Lao, J.; Madani, J.; Puértolas, T.; Álvarez, M.; Hernández, A.; Pazo-Cid, R.; Artal, Á.; Antón Torres, A. Liposomal doxorubicin in the treatment of breast cancer patients: a review. Journal of Drug Delivery 2013, 2013, 2090–3014.

(38) Tacar, O.; Sriamornsak, P.; Dass, C. R. Doxorubicin: an update on anticancer molecular action, toxicity and novel drug delivery systems. Journal of Pharmacy and Pharmacology 2013, 65 (2), 157–170.

(39) El-Sayed, A.; Masuda, T.; Akita, H.; Harashima, H. Stearylated INF7 peptide enhances endosomal escape and gene expression of PEGylated nanoparticles both in vitro and in vivo. Journal of pharmaceutical sciences 2012, 101 (2), 879–882.

(40) Nakase, I.; Kobayashi, S.; Futaki, S. Endosome-disruptive peptides for improving cytosolic delivery of bioactive macromolecules. Peptide Science 2010, 94 (6), 763–770.

(41) Kaksonen, M.; Roux, A. Mechanisms of clathrin-mediated endocytosis. Nature reviews Molecular cell biology 2018, 19 (5), 313–326.

(42) Feng, Y.; Press, B.; Wandinger-Ness, A. Rab 7: an important regulator of late endocytic membrane traffic. The Journal of cell biology 1995, 131 (6), 1435–1452.

